# Unravelling the complexity of silk sericins: *P150/sericin 6* is a new silk gene in *Bombyx mori*

**DOI:** 10.1101/2023.09.22.558982

**Authors:** Bulah Chia-hsiang Wu, Valeriya Zabelina, Martina Zurovcova, Michal Žurovec

## Abstract

Sericins are a small family of highly divergent proteins that serve as adhesives and coatings for silk fibers and are produced in the middle part of the silk gland. So far, five genes encoding sericin proteins have been found in *Bombyx mori*. Sericins 1 and 3 are responsible for silk adhesion in the cocoon, while sericins 2, 4, and 5 are present in non-cocoon spun silk of younger larvae (including the early last instar). We found a new gene, which we named *P150/sericin 6*, which appears to be an ortholog of the sericin-like protein previously found in *Galleria mellonella*. The *B. mori* sequence of the *P150/sericin 6* ORF was previously incorrectly predicted and assigned to two smaller, uncharacterized genes. We present a new *P150/sericin 6* gene model and show that it encodes a large protein of 467 kDa. It is characterized by repeats with a high proportion of threonine residues and a short conserved region with a cysteine knot motif (CXCXCX) at the C-terminus. Expression analysis has shown that *B. mori P150/ser6* has low transcriptional level in contrast to its *G. mellonella* homolog. We also discuss the synteny of homologous genes on corresponding chromosomes between moth species and possible phylogenetic relationships between *P150/ser6* and cysteine knot mucins. Our results improve our understanding of the evolutionary relationships between adhesion proteins in different lepidopteran species.

## Introduction

Silk is a secretory product of several arthropod groups, including insects with the best known being produced by specialized larval salivary glands (the silk glands, SG) of the silkworm. Silk fibers are composed mainly of two types of proteins: fibroins and sericins. Fibroins are the central structural proteins that give silk strength and durability. In most moth species, fibroin is a complex of three protein subunits, fibroin heavy chain (Fib-H), fibroin light chain (Fib-L), and fibrohexamerin/P25 (Fhx/P25)[1-3]. They are produced in the posterior part of SG [4]. Sericins, on the other hand, are coating proteins produced in several layers in the middle part of the SG [5]. Sericins are a small highly divergent family of adhesives that help to glue the fibers together so that silk can form intricate structures such as cocoons or feeding tubes. The use of sensitive proteomic methods and the sequencing of transcriptomes and genomes of individual species indicate that the family of sericin proteins is larger than previously thought. The large divergence among sericin proteins suggests that they have evolved to perform slightly different functions depending on the specific needs of the silk-producing organism.

Two major sericin genes of *B. mori* have been found to produce cocoon sericins: Sericin 1 (*Ser1*) and Sericin 3 (Ser3) [6, 7]. Silkworm mutants carrying a truncated *ser1* gene failed to spin or produced coarse cocoons, suggesting that it is involved in reducing friction during spinning [8]. The product of another sericin gene, sericin 2 (Ser2), and the products of two recently identified genes sericin 4 (Ser4) and sericin 5 (Ser5), have been described in non-cocoon silk and are spun by younger larvae (including early last instar larvae) [9-12]. The presence of Ser2, Ser4, and Ser5 in non-cocoon silks suggests that they may have a specific function during these developmental stages. In this study, we describe another adhesive protein from *B. mori* silk that is homologous to a previously identified sericin-like protein, P150, found in the larval cocoons of *G. mellonella* and *Ephestia kuehniella* (superfamily Pyraloidea) [13, 14].

We show here that there is at least one additional sericin-like protein in the silk of *B. mori*, which we named P150/Ser6 and which is similar to the previously discovered sericin protein P150 of *G. mellonella* [13]. Our results suggest that *B. mori* sequences similar to P150 were misannotated in both GenBank and Silk Base and assigned to two different genes with unknown functions. We present a new model of the *P150/ser6* gene and show that it is a large gene with 4 exons expressed in the middle part of the SG. The *P150/ser6* protein product is found in the pre-cocoon silk, and to a lesser extent in the inner cocoon layer formed at the end of the last instar. The newly discovered protein adds to the existing sericin family of the silkworm. Our results improve our understanding of the functions of silk proteins and the evolutionary relationships among adhesion proteins in different lepidopteran species.

## 2. Materials and methods

### 2.1. Silkworm strains and datasets used

A non-diapausing *B. mori* strain, *w1-pnd* (*white egg 1, non-pigmented and non-diapausing egg*), was used in the experiments as a wild type (*wt*) strain. Larvae were reared on mulberry leaves at 25°C. A list of RNA-seq datasets from NCBI Sequence Read Archive (SRA) is provided in Table S1. (Supplementary Table S1).

### 2.2. RNA extraction and RT-PCR

To verify the structure of *P150/ser6*, total RNA was extracted from the SGs from SGs of 3-5-day-old fifth-instar larvae using Trizol reagent (Invitrogen, Carlsbad, CA, USA) according to the manufacturer’s instructions. The first cDNA strand was synthesized using 0.5 μg of total RNA as template. The cDNA product was used to verify the last exon junction of B. mori P150/Ser6 and its expression level in different tissues. Primers were designed using the Geneious Prime software platform (Biomatters, Auckland, New Zealand; version 2021.2.2) and are listed in Supplementary Table S2.

qPCR was performed using HOT FIREPol EvaGreen qPCR Mix Plus (Solis BioDyne, Tartu, Estonia). The PCR reaction volume of 20 µl contained 5 µl diluted cDNA and 250 nM primer. Amplification was performed using a Rotor-Gene Q MDx 2plex HRM (Qiagen, Hilden, Germany) for 45 cycles (95°C for 15 s; annealing temperature matched to the primer pair for 30 s; 72°C for 20 s) after an initial denaturation step (95°C for 15 min). Each sample was analyzed in triplicate. Results were analyzed using Rotor Gene Q software (version 2.3.5). Elongation factor 1 alpha (EF1a, NM _001044045.1) was used as a reference gene, and the relative expression of target genes was calculated using the 2−ΔΔCT method [15]. Statistical analysis was performed using Student’s t-test in R (version 4.1.1); p values < 0.05 were considered statistically significant. Detailed statistical analysis is provided in Supplementary Table S3.

### 2.3. Proteomic analysis and data mining

Protein analysis of *B. mori* cocoons and database searches were performed at the Proteomics Core Facility (BIOCEV, Vestec, Czech Republic) as previously described [16]. Approximately 10 mg of the silk cocoon was boiled in 8 M urea, and samples were further processed using solid-phase enhanced sample preparation technology (SP3 beads) [17]. Samples were then digested with trypsin, and the resulting peptides obtained were subjected to liquid chromatography - MS. Four *wt* cocoons were analyzed in parallel. In addition, raw proteomic data deposited in public databases [18] on the composition of individual silk layers in the *B. mori* cocoon were reanalyzed. The obtained MS/MS spectra were matched against the UNIPROT database (hwww.uniprot.org), which was enriched for the newly discovered P150/Ser6. Quantification was performed using label-free algorithms, and data were analyzed using MaxQuant and Perseus v.1.5.2.4 [19, 20].

### 2.4. Chromosomal localization and collinearity analysis

The genome assemblies and annotated information of *B. mori* (GCF_014905235.1), *E. kuehniella* [14, 21], and *G. mellonella* (GCF_026898425.1) were processed and submitted to the GENESPACE software [22] for syntenic analysis. Plots showing the microsyntenic relationships were then generated based on the best mutual hits between the three species and visualized using the R package ggplot2 [23].

### 2.5. Phylogenetic analysis

Codon-based alignment of the *P150/ser6* and cysteine knot mucin 3’ ends was performed using MEGA7 software following the MUSCLE method [24]. The phylogram was generated using the IQ-TREE server [25, 26], which included both the selection of the best substitution model by ModelFinder [27] and tree inference using MLE (ultrafast bootstrap, 1,000 replicates).

## 3. Results

### 3.1. Identification of P150/ser6 gene in B. mori

To identify the homolog of the major sericin gene P150 described previously in *G. mellonella* and *E. kuehniella*, we performed a BLAST search against the *B. mori* genome. We found two adjacent homologous regions in the genomic sequence, which were predicted to be parts of two *B. mori* genes. Most of the homologous sequence belonged to *XP_037870317*.*1*; the remaining C-terminus was predicted to be a separate gene *XP_037870336*.*1*. The *B. mori* sequence shared 47.5% identity per 70 C-terminal amino acid residues with *G. mellonella P150*. We hypothesized that the predicted gene models were incorrect and that the sequences of both *B. mori* genes were part of one large *P150/ser6* gene. To test that the two putative *B. mori* genes represent a single gene and produce a single large mRNA, we designed RT-PCR primers that fuse the last two exons of the predicted *XP_037870317*.*1* with the second exon *XP_037870336*.*1* and amplified a fragment that confirmed the integrity of the putative large exon *XP_037870317*.*1* (Fig. 1A). We also designed primers that bridge both exons of *XP_037870336*.*1*. As shown in Figure 1B, the amplified cDNA fragments supported the hypothesis of a single large *P150/ser6* gene.

**Fig. 1.**
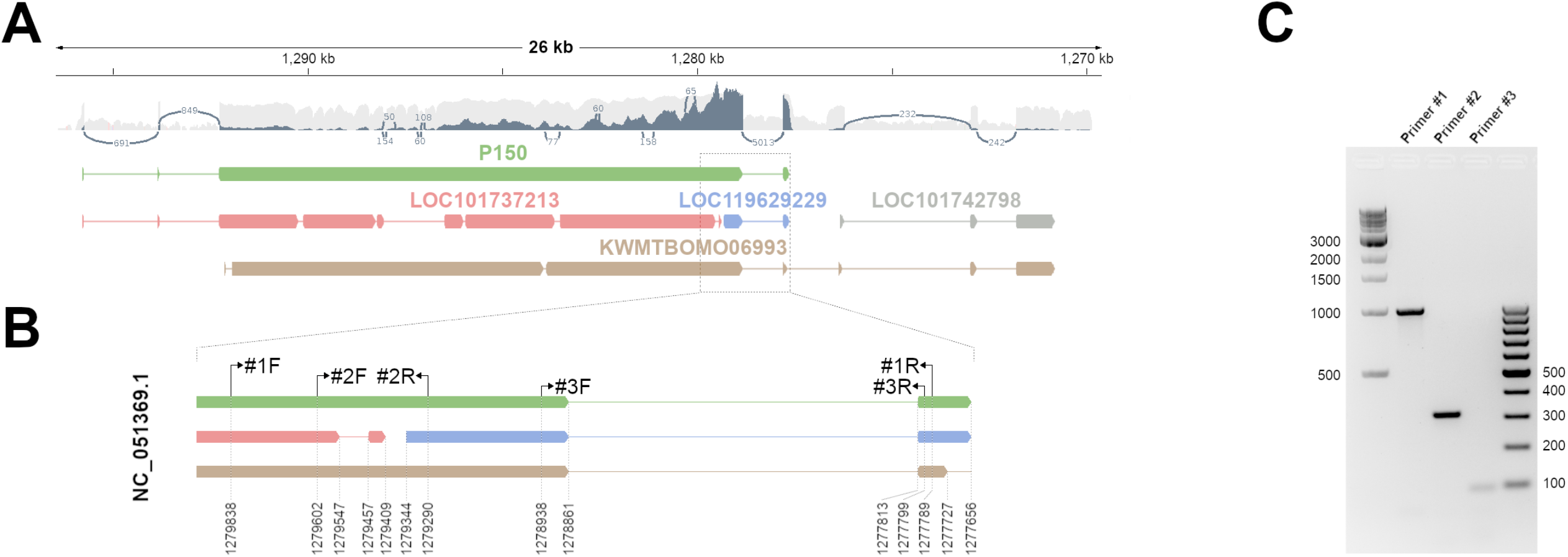
Different models of a *B. mori* genomic region containing sequences similar to the P150 protein of *G mellonella*. (**A**) The genomic region is part of chromosome 12 (NC_051369.1: 1270000-1280000, strand flipped). The exon-intron structure of the *P150/ser6* gene, consisting of four exons and three introns, as predicted by our analysis, is shown in green. Below - comparison with current models in the NCBI database the following gene models are shown: *XP_037870317*.*1* (pink); *XP_037870336*.*1* (blue); *XP_004929407*.*1* (gray); and the model of the gene *KWMTBOMO06993* in Silkbase (brown). The size bar in kilobase pairs is shown at the top. **(B)** The lower inset *shows* an approximately 2.2-kb region with the chromosomal coordinates and positions of the primer pairs used in this study. (**C**) Verification of the 3’ region of the newly designed *P150/ser6* gene model by RT-PCR and agarose gel electrophoresis. The expected product sizes of primer pairs #1F-1R, #2F-2R, and #3F-3R were 1003 bp, 313 bp, and 93 bp, respectively. The electrophoretogram contains a 1 kb ladder (lane 1) and 100 bp ladder (lane 5).

The new gene model of *P150/ser6* is shown in Figure 1A. The entire gene spans approximately 20 kb and consists of four exons and three introns (Fig. 1A). The first two exons of *P150/ser6* encode a signal peptide and part of the short N-terminal nonrepetitive sequence. The third exon is very large (containing 94% of the ORF) and contains two central repetitive regions flanked by unique sequences. The last exon contains a short ORF and a stop codon. The entire gene contains an ORF encoding 4552 amino acids including a signal peptide (19 amino acid residues).

### 3.2. Putative P150/Ser6 protein

The deduced protein product of the P150/ser6 gene is a large protein of 467 kDa consisting of 4552 amino acid residues. It contains a signal peptide of 19 amino acid residues in length, followed by a central portion consisting of a 616 amino acid non-repetitive region, followed by 45 copies of a 30-amino acid repeat 1, a 34-amino acid non-repetitive linker, a repeat 2 consisting of 73 copies of 35 amino acids, and a nonrepetitive C-terminus of 388 amino acids (the complete sequence is shown in Text. S1). As shown in shown in Fig. 2, P150/Ser6 consists of two types of highly conserved, threonine -rich repeat blocks (Supplementary Table S4). The P150/Ser6 protein is relatively highly hydrophilic (hydropathy index = -0.672 compared to -1.118 of Ser1). The B. mori P150/Ser6 contains more than 27% threonine, 14% serine, and 12% alanine residues. The C-terminus (encoded by the last exon) contains a short, conserved cysteine knot motif (Supplementary Table S5).

**Fig. 2.**
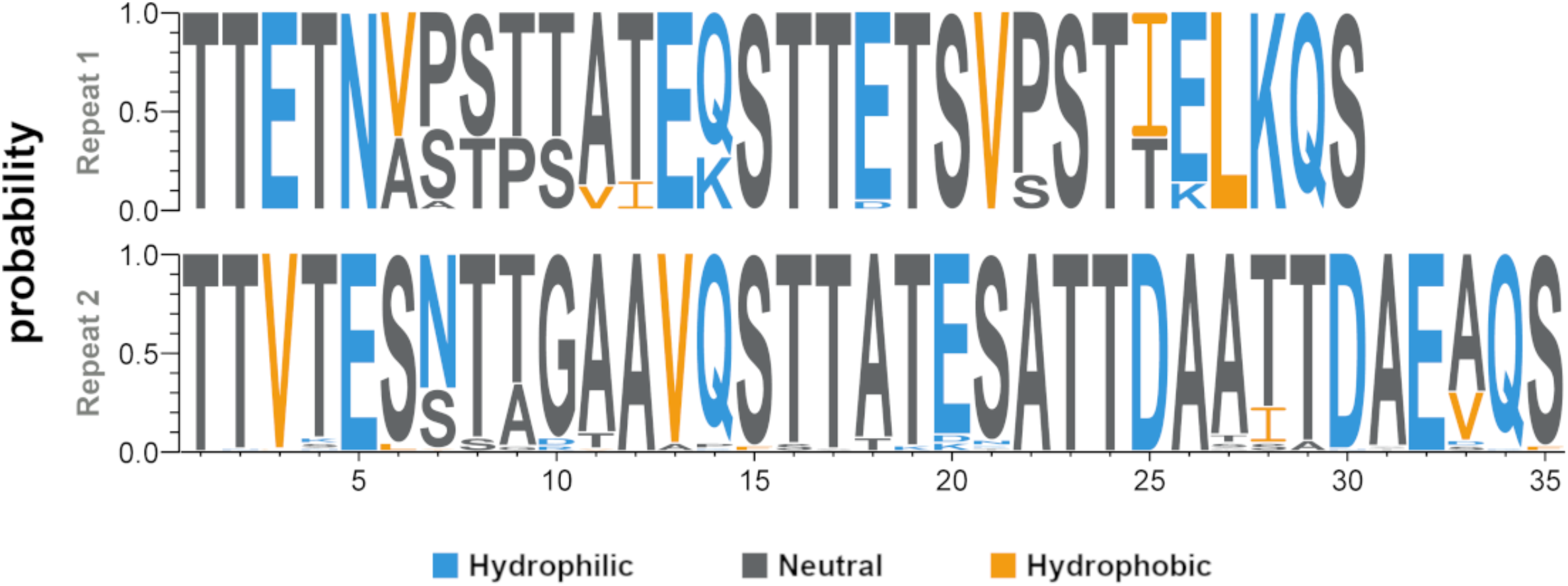
Amino acid sequence logos. The sequence logos show the conservation of amino acids in two types of repeats found in the *B. mori* P150/Ser6 protein. Hydrophobicity of an amino acid is indicated by color: hydrophilic (blue; RKDENQ); neutral (dark gray; SGHTAP); hydrophobic (orange; YVMCLFIW).

Compared to the P150 proteins of *G. mellonella* and *E. kuehniella*, the *B. mori* P150/Ser6 is almost three times larger, less hydrophilic, and contains fewer serine residues. There is a very little conservation between the P150/Ser6 proteins except for the C-terminal amino acids (Supplementary Table S6).

### 3.3. P150/ser6 mRNA is specifically expressed in MSG

To determine whether the *B. mori* homolog of P150/Ser6 is specifically expressed in silk glands, we isolated mRNAs from different parts of the silk glands and from control tissues (intestine and ovary) of day 3-5 last instar larvae, prepared cDNA, and performed qPCR. As shown in Figure 3, P150/Ser6 mRNA is highly specific for MSG and ASG, whereas it is absent in PSG and control tissues.

**Fig. 3.**
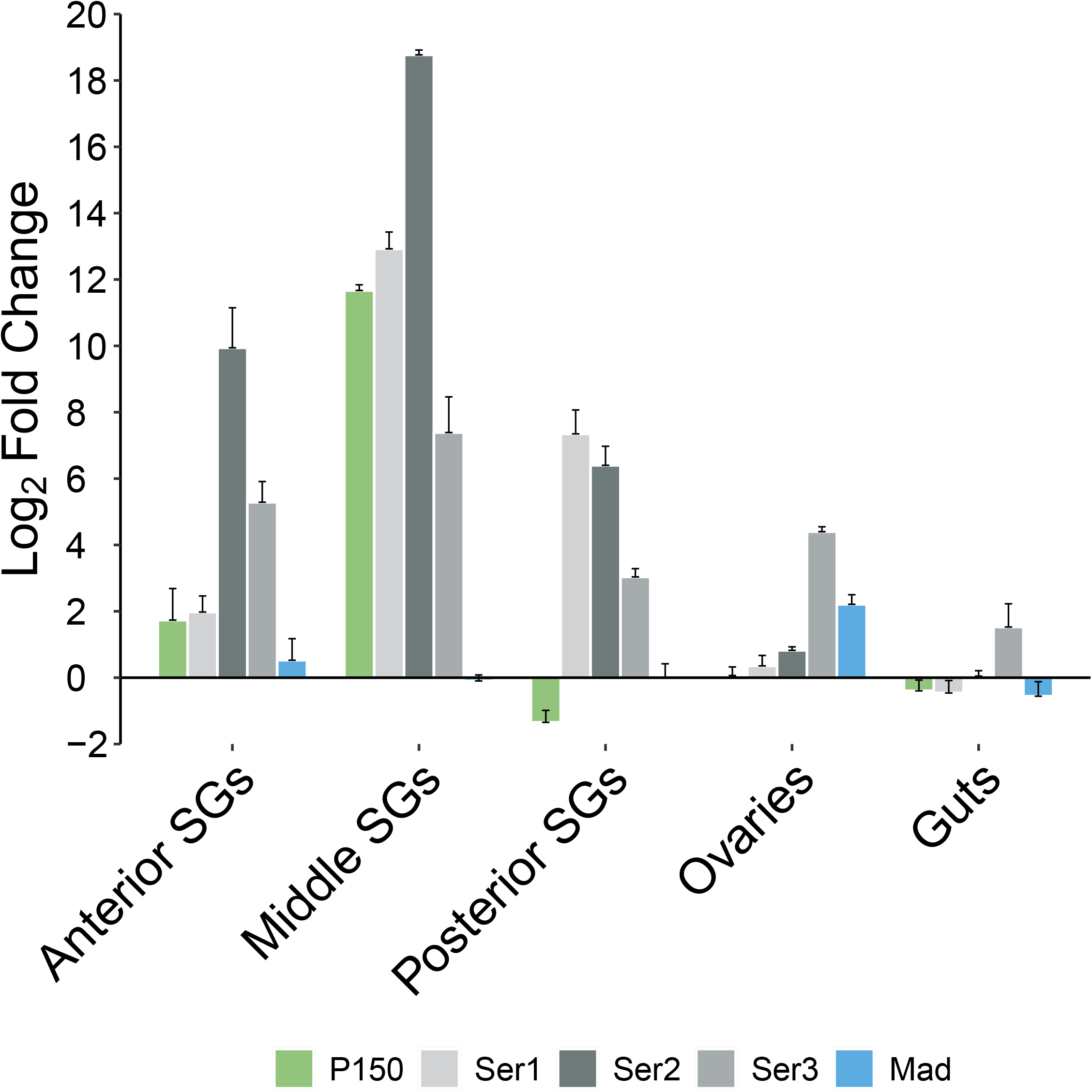
Quantitative PCR (qPCR) analysis of gene expression of silk gland-specific genes (*ser1, ser2, ser3*, and *P150/ser6*) and control (non-silk gland-specific gene MAD) in different tissues. Statistical differences were evaluated using Student’s t-test (see Supplementary data); error bars are SD.

In addition, we reanalyzed the publicly available RNAseq data for silk gene expression from previous experiments [18, 28]. We used a new annotation of *P150/ser6* and quantified transcript abundance using Kallisto software, and estimated fold changes using DESeq2 (see material and methods). As shown in Figure S1, the results support our data above and indicate that *P150/ser6* mRNA is highly specific to MSG and its maximal expression occurs in the middle part of MSG, similar to that of Ser2 and Muc-12. In addition, the maximal expression of Ser1 and Ser3 is found in the middle part of MSG, with a low expression level also in PSG (Fig. S1). The presence of sericin mRNAs in the posterior SG sample may be caused by different separation sites between the posterior and middle SG during tissue dissection.

### 3.4. Quantitative proteomic analysis of silk samples

To test whether the P150/Ser6 protein is present in *B. mori* cocoons, we performed MS proteomic analyses of silk from wt cocoons. The results were examined using the Andromeda search engine integrated into the MaxQuant software. The relative abundance of proteins was determined by label-free quantification. We identified 118 proteins using the false discovery rate (FDR) of 1% for protein identification. The intensity of each protein in the biological samples was in strong agreement. A summary of the identified proteins from triplicate analysis of each cocoon is shown in Table 1.

The results of the proteomic analysis show that the expression of P150/Ser6 protein is quite low, similar to those of Ser2, and Muc-12, which are also present at low intensities at the instrumental detection limit (IDL). In contrast, the data (Fig. 4A) showed that Ser1 and Ser3 are the most abundant components of the cocoon silk with concentrations at least five orders of magnitude higher than P150/Ser6 (Fig. 4A).

**Fig. 4.**
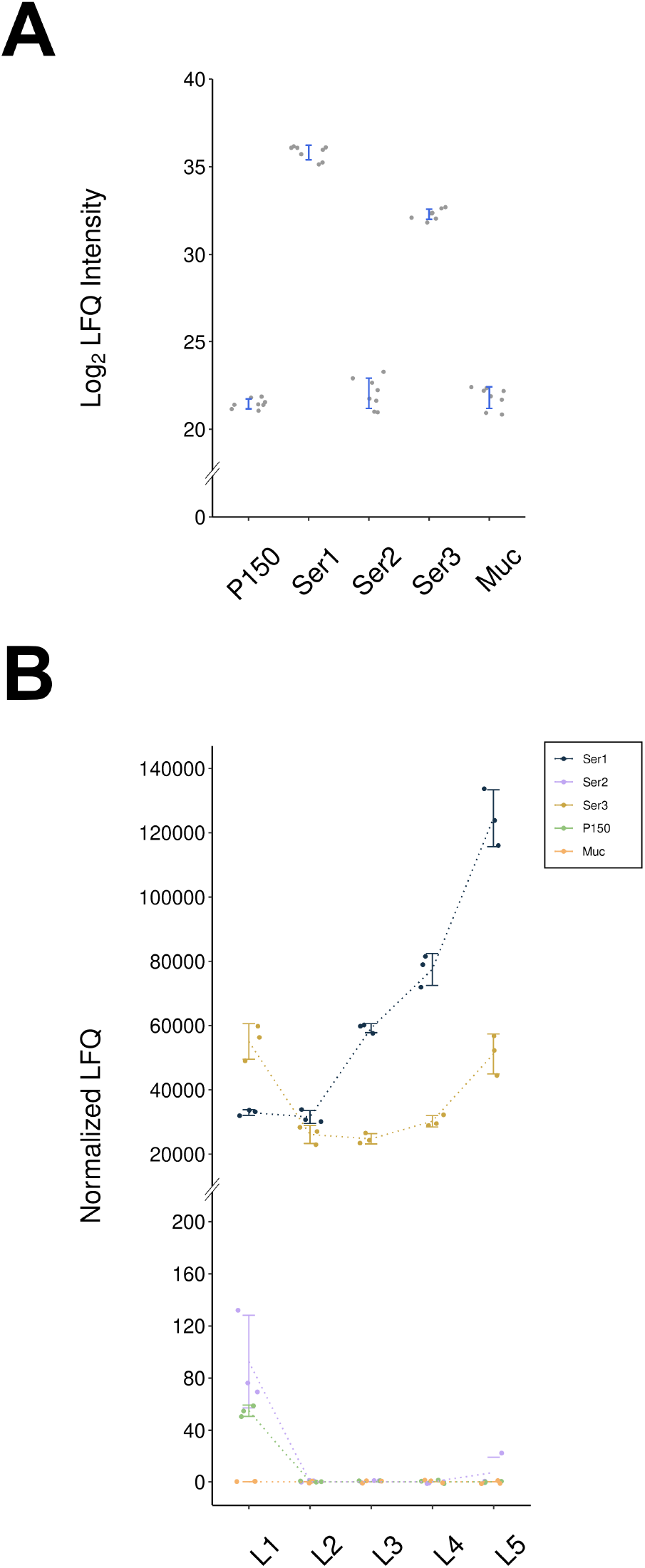
Proteomic analysis of *B. mori* silk proteins (**A**) from *wt* cocoons as previously described [16]; and (B) individual cocoon layer data from a public repository [18]. Label-free quantification (LFQ) of silk proteins from cocoons was calculated using MaxQuant. LFQ intensities were log2-transformed. Relative protein contents in cocoon silk were analyzed using MaxQuant/Andromeda (eight experiments). Error bars indicate the standard deviation. Proteomic analysis confirmed that Ser1 and Ser3 were the most abundant silk components. The other proteins, including P150/ser6, Ser2, and Muc-12, were present at low levels at the instrumental detection limit (IDL).

To determine the protein abundance in the different cocoon layers and whether P150/Ser6, Muc-12, and Ser2 are coordinately expressed, we also re-analyzed the existing proteomics data in the public repository [18] using our new annotation of P150/Ser6. The abundance of P150/Ser6, Muc-12 and Ser2 in cocoons is shown in Figure 4B. All three proteins are found at very low levels, with P150/Ser6 and Ser2 being the most abundant in the innermost cocoon layer (layer 1), whereas Muc-12 is found in the outermost layer. Taken together, our data show that the abundance of P150/Ser6 and Muc-12 in cocoons is low and differs in localization and timing of secretion, with the outermost layer being secreted first, whereas the innermost layer is produced at the end of spinning.

### 3.5. Synteny in regions coding for P150/ser6 genes in Lepidoptera

A previous study on the pyralid moths *G. mellonella* and *E. kuehniella* showed that the known sericin genes, with the exception of P150/ser6, lie within the cluster of orthologous genes in the corresponding chromosomal regions [14]. Such microsynteny can be also observed between *B. mori* and *G. mellonella* or *E. kuehniella* (Fig 5). The results also show a number of local rearrangements and duplications in this region including the expansion of several sericin genes in *G. mellonella* compared to related moths [14].

**Fig. 5.**
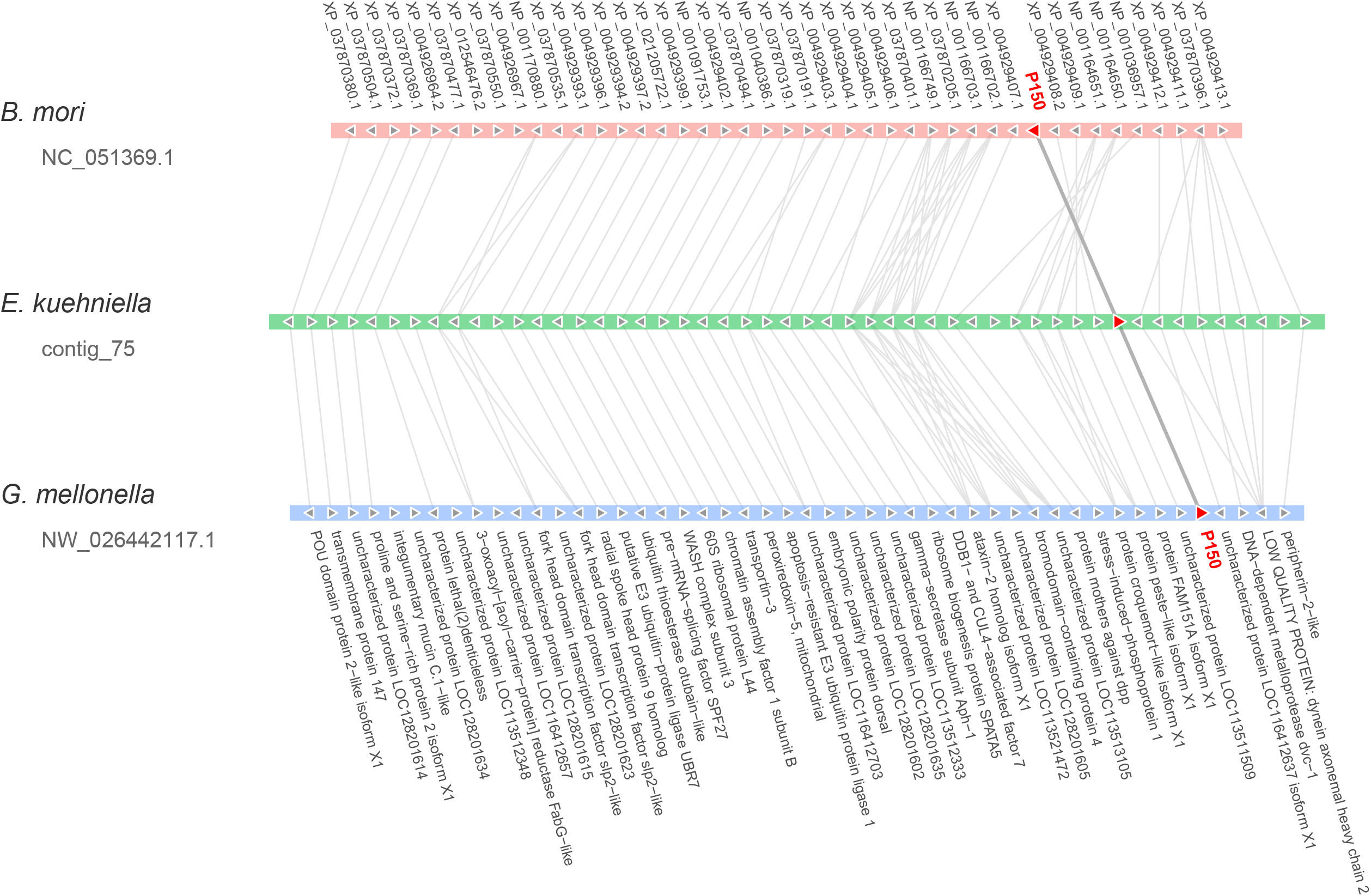
Microsynteny maps of *P150/ser6* and their flanking genes. Horizontal color blocks indicate chromosomal segments in each species. Homologous genes and gene orientation are represented by left- and right-pointing triangles and connected by lines. *Detected P150/ser6* homologs are shown in red.

In contrast, the *P150/ser6* gene is located on a different chromosome in a more conserved region of a separate cluster, between the genes encoding *metalloprotease 1* and *croquemont 1*. As shown in Figure 5, the region on chromosome 12 of *B. mori* has a well-conserved synteny, except for an inversion that places the *P150/ser6* region in the opposite direction to the surrounding genes.

### 3.6. P150/Ser6 may be related to Muc-12

To investigate the evolution of P150/ser6, we searched for homologous sequences in insect genomes using BLAST and the C-terminal conserved sequence as bait. We found no obvious orthologs in non-lepidopteran insects, suggesting that the *P150/ser6* gene appears to be specific to Lepidoptera. We also found no P150/Ser6 ortholog in members of the superfamily Papilionoidea.

P150/Ser6 proteins are highly divergent, and homologous proteins from different Lepidopteran families are difficult to align, except for the conserved C-terminus (the consensus sequence is shown in Fig. 3). The most prominent conserved motif is the cysteine knot (CXCXCX) sequence, which is located 12–29 amino acid residues away from the C-terminus.

Interestingly, there are other silk gland-specific proteins, which also contain cysteine knot sequences. One of these is Mucin-12, which is reminiscent of P150/Ser6 because of its size and the repetitive structure of its molecule. To learn more about the relationship between cysteine knot mucins and P150/Ser6 sequences, we constructed a dendrogram of P150/Ser6 and Muc-12 C-termini from representatives of several lepidopteran families (Fig. 6). The resulting phylogenetic tree is robust and distinguishes both clades—P150/Ser6 and Muc-12—with high support except for the sequences from the most primitive species (Fig. 6). The sequence alignment and consensus are shown in Figure 6B.

**Fig. 6.**
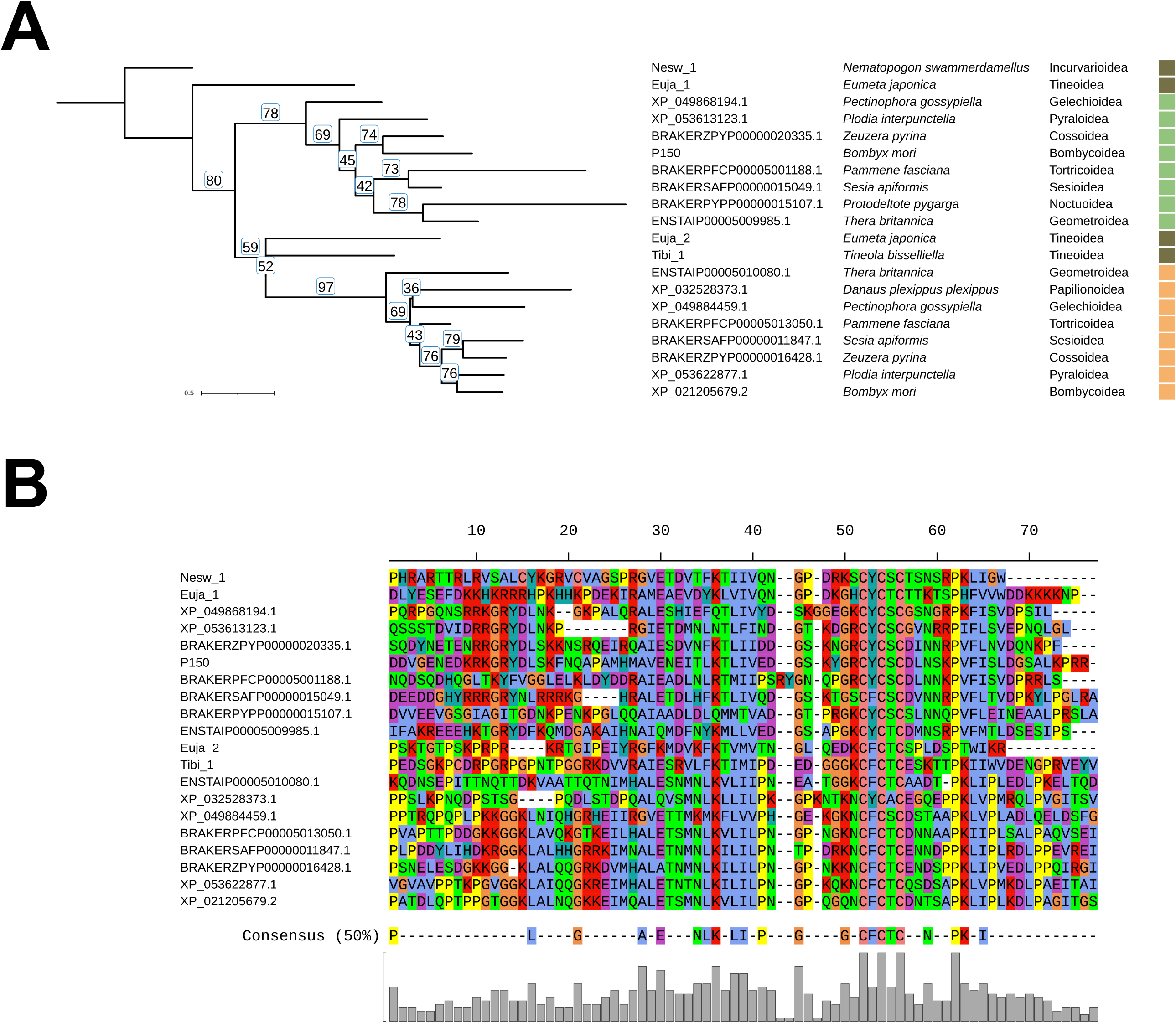
Relationship between P150/Ser6 and cysteine knot mucin sequences. (**A**) Maximum likelihood phylogenetic tree based on the alignment of the C-terminal amino acid sequences of P150/Ser6 and Muc-12 homologs in selected lepidopteran species. The *Nesw_1* transcript from *N. swammerdamellus*, (Incurvarioidea), the most primitive in this group, was selected for tree rooting. See Supplementary Text S2 and Supplementary Table S7 for sequences. (**B**) Alignment of the C-terminal regions of P150/ser6 proteins from representative lepidopteran species. Sequences include the characteristic CXCXCX region, which is well conserved between species.

## 4. Discussion

We discovered a new sericin-like gene *P150/ser6* in the genome of *B. mori*, based on homology with a similar gene in *G. mellonella*. We also found that the region on chromosome 12 where this gene is located was not correctly annotated. Previous gene models of *P150/ser6* contained a homologous sequence that was split into two putative *B. mori* genes. Here, we present the correct *P150/ser6* gene model and show that its ORF encodes a large protein with a repetitive structure that is specifically expressed in SG.

Structure prediction of large genes is still a problem [29]. The best results are obtained by comparing genomic and cDNA sequences or by using proteomic data [30]. Previous identification of *P150/ser6* sequences in *G. mellonella* and *E. kuehniella* was successfully performed using this approach [13, 14]. In contrast, finding the *B. mori P150/ser6* homolog has been more difficult due to its relatively low expression. Here we show that *P150/ser6* and possibly some other silk genes can be identified on the basis of the homology of a sufficiently conserved, relatively short motif. The identification of *P150/ser6* was supported by microsynteny between the corresponding genomic regions in *B. mori* and *G. mellonella* (Fig. 5).

Previous results showed that the homolog of P150/Ser6 is one of the most abundant silk proteins in *G. mellonella* [13]. In contrast, our results show that it is present only at trace levels in *B. mori* silk. The P150/Ser6 protein product of *B. mori* is found mainly in the inner cocoon layer formed at the end of the last instar and also in the non-cocoon silk from earlier instar larvae, as suggested by the data of Dong et al. [28]. Consistently, Ser2, Ser4 and Ser5 are present in the non-cocoon silk [9-11]. Non-cocoon silk has previously been associated with the initial stage of silk spinning and is responsible for holding the molting larva and fixing the cocoon to a suitable substrate [11]. Alternatively, the sericin sequences of *B. mori* sericin sequences could have high serine content (Supplementary Table S5), as only simple sericins are digestible by cocoonase [31]. Sericins containing less serine would be restricted to non-cocoon silk. Mutants in cocoonase result in adults trapped in the cocoon.

Why are there multiple sericins in moth silks? Unlike spiders, moths have only one fibroin gene, which is considered to be the main structural component of silk. However, silk is modified differently in different moth species, and sericins appear to be the most variable components, important for building three-dimensional silk structures and contributing to silk strength and toughness [32]. For example, *G. mellonella* builds very dense feeding tubes and cocoons that are needed for larval protection in hives. Sericins and other soluble silk components make up about 48% of the cocoon mass of *G. mellonella*, whereas *B. mori* cocoons contain only about 26% soluble cocoon proteins. Some moths, including those of *Samia ricini* contain as little as 16 % soluble proteins [11, 33]. The number of sericin genes can also vary widely, for example, *G. mellonella* contains twice as many sericin genes as *E. kuehniella* [14].

The cysteine knot motif (CXCXCX) encoded by the C-termini of some SG-specific genes is similar to the motif described in some mammalian growth factors, including the VEGF family [34]. The function of cysteine knot motifs has been suggested for protein structural integrity. In addition to P150/Ser6 and Muc-12, there are at least two other *B. mori* SG-specific proteins that carry this motif. One of these, a putative 53 kDa non-repetitive protein, is also expressed in silk glands and has homologs in other moths and even caddisflies. The other protein, designated egalitarian protein homolog (LOC101741849), is more conserved and appears to be well separated from other cysteine knot proteins [35].

The general similarity of P150/Ser6 and Muc-12 suggests that they may have a common origin. P150/Ser6 and Muc-12 are both large, highly divergent proteins. They have repetitive sequences encoding ORFs with simple amino acids and both are likely to serve as silk adhesives [13]. Our phylogram (Fig. 6) distinguishes the two clades P150/Ser6 and Muc-12 with good support, except for the sequences in the most primitive species, where it is difficult to place them confidently in the phylogram. The evolutionary relationship between P150/Ser6 and mucins with a cysteine node motif is not clear. The question of whether P150/Ser6 and cysteine knot mucins share a common ancestor and have diverged extensively or whether they are the product of convergent evolution remains an important question for future research.

## Author contributions

MiZ: supervision writing; BChW: investigation, data analysis; writing; MaZ: phylogenetic analysis; VZ: *B. mori* rearing and staging.

## Data and materials availability

Data will be made available on request.

## Acknowledgments

This research was supported by European Community’s Program Interreg Bayern – Tschechien BYCZ01-039. This publication is also supported by the project “BIOCEV – Biotechnology and Biomedicine Centre of the Academy of Sciences and Charles University” (CZ.1.05/1.1.00/02.0109), from the European Regional Development Fund.

## Supplementary materials

## References

[1] K. Tanaka, K. Mori, S. Mizuno, Immunological Identification of the Major Disulfide-Linked Light Component of Silk Fibroin, J Biochem-Tokyo 114(1) (1993) 1–4.

[2] M. Zurovec, D. Kodrik, C. Yang, F. Sehnal, K. Scheller, The P25 component of Galleria silk, Mol Gen Genet 257(3) (1998) 264–70.

[3] M. Zurovec, M. Vaskova, D. Kodrik, F. Sehnal, A.K. Kumaran, Light-chain fibroin of Galleria mellonella L, Mol Gen Genet 247(1) (1995) 1–6.

[4] F. Lucas, K.M. Rudall, Extracellular fibrous proteins: The silks., in: M. Florkin, E.H. Stotz (Eds.), Comprehensive Biochemistry, Elsevier, Amsterdam, 1968, pp. 475–558.

[5] A. Shibukawa, Studies on the silk substance within the silk gland in the silkworm, Bombyx mori L., Bull. Sericult. Exp. Sta. 15 (1959) 401.

[6] H. Okamoto, E. Ishikawa, Y. Suzuki, Structural analysis of sericin genes. Homologies with fibroin gene in the 5’ flanking nucleotide sequences, J Biol Chem 257(24) (1982) 15192–9.

[7] Y. Takasu, H. Yamada, T. Tamura, H. Sezutsu, K. Mita, K. Tsubouchi, Identification and characterization of a novel sericin gene expressed in the anterior middle silk gland of the silkworm Bombyx mori, Insect Biochem Mol Biol 37(11) (2007) 1234–40.

[8] Y. Takasu, T. Iizuka, Q. Zhang, H. Sezutsu, Modified cocoon sericin proteins produced by truncated Bombyx Ser1 gene, The Journal of Silk Science and Technology of Japan 25 (2017) 35–47.

[9] Z.M. Dong, K.Y. Guo, X.L. Zhang, T. Zhang, Y. Zhang, S.Y. Ma, H.P. Chang, M.Y. Tang, L.N. An, Q.Y. Xia, P. Zhao, Identification of Bombyx mori sericin 4 protein as a new biological adhesive, Int J Biol Macromol 132 (2019) 1121–1130.

[10] K. Guo, X. Zhang, D. Zhao, L. Qin, W. Jiang, W. Hu, X. Liu, Q. Xia, Z. Dong, P. Zhao, Identification and characterization of sericin5 reveals non-cocoon silk sericin components with high beta-sheet content and adhesive strength, Acta Biomater 150 (2022) 96–110.

[11] B. Kludkiewicz, Y. Takasu, R. Fedic, T. Tamura, F. Sehnal, M. Zurovec, Structure and expression of the silk adhesive protein Ser2 in Bombyx mori, Insect Biochem Molec 39(12) (2009) 938–946.

[12] J.J. Michaille, A. Garel, J.C. Prudhomme, Cloning and Characterization of the Highly Polymorphic Ser2 Gene of Bombyx-Mori, Gene 86(2) (1990) 177–184.

[13] B. Kludkiewicz, L. Kucerova, T. Konikova, H. Strnad, M. Hradilova, A. Zaloudikova, H. Sehadova, P. Konik, F. Sehnal, M. Zurovec, The expansion of genes encoding soluble silk components in the greater wax moth, Galleria mellonella, Insect Biochem Mol Biol 106 (2019) 28–38.

[14] B.C. Wu, I. Sauman, H.O. Maaroufi, A. Zaloudikova, M. Zurovcova, B. Kludkiewicz, M. Hradilova, M. Zurovec, Characterization of silk genes in Ephestia kuehniella and Galleria mellonella revealed duplication of sericin genes and highly divergent sequences encoding fibroin heavy chains, Front Mol Biosci 9 (2022) 1023381.

[15] K.J. Livak, T.D. Schmittgen, Analysis of relative gene expression data using real-time quantitative PCR and the 2(-Delta Delta C(T)) Method, Methods 25(4) (2001) 402–8.

[16] V. Zabelina, Y. Takasu, H. Sehadova, N. Yonemura, K. Nakajima, H. Sezutsu, M. Sery, M. Zurovec, F. Sehnal, T. Tamura, Mutation in Bombyx mori fibrohexamerin (P25) gene causes reorganization of rough endoplasmic reticulum in posterior silk gland cells and alters morphology of fibroin secretory globules in the silk gland lumen, Insect Biochem Mol Biol 135 (2021) 103607.

[17] C.S. Hughes, S. Moggridge, T. Muller, P.H. Sorensen, G.B. Morin, J. Krijgsveld, Single-pot, solid-phase-enhanced sample preparation for proteomics experiments, Nat Protoc 14(1) (2019) 68–85.

[18] Y. Zhang, P. Zhao, Z. Dong, D. Wang, P. Guo, X. Guo, Q. Song, W. Zhang, Q. Xia, Comparative proteome analysis of multi-layer cocoon of the silkworm, Bombyx mori, PLoS One 10(4) (2015) e0123403.

[19] J. Cox, M.Y. Hein, C.A. Luber, I. Paron, N. Nagaraj, M. Mann, Accurate proteome-wide label-free quantification by delayed normalization and maximal peptide ratio extraction, termed MaxLFQ, Mol Cell Proteomics 13(9) (2014) 2513–26.

[20] S. Tyanova, T. Temu, P. Sinitcyn, A. Carlson, M.Y. Hein, T. Geiger, M. Mann, J. Cox, The Perseus computational platform for comprehensive analysis of (prote)omics data, Nat Methods 13(9) (2016) 731–40.

[21] S. Visser, A. Volenikova, P. Nguyen, E.C. Verhulst, F. Marec, A conserved role of the duplicated Masculinizer gene in sex determination of the Mediterranean flour moth, Ephestia kuehniella, PLoS Genet 17(8) (2021) e1009420.

[22] J.T. Lovell, A. Sreedasyam, M.E. Schranz, M. Wilson, J.W. Carlson, A. Harkess, D. Emms, D.M. Goodstein, J. Schmutz, GENESPACE tracks regions of interest and gene copy number variation across multiple genomes, Elife 11 (2022).

[23] H. Wickham, ggplot2: Elegant Graphics for Data Analysis, Use R (2009) 1–212.

[24] S. Kumar, G. Stecher, K. Tamura, MEGA7: Molecular Evolutionary Genetics Analysis Version 7.0 for Bigger Datasets, Mol Biol Evol 33(7) (2016) 1870–4.

[25] D.T. Hoang, O. Chernomor, A. von Haeseler, B.Q. Minh, L.S. Vinh, UFBoot2: Improving the Ultrafast Bootstrap Approximation, Mol Biol Evol 35(2) (2018) 518–522.

[26] L.T. Nguyen, H.A. Schmidt, A. von Haeseler, B.Q. Minh, IQ-TREE: a fast and effective stochastic algorithm for estimating maximum-likelihood phylogenies, Mol Biol Evol 32(1) (2015) 268–74.

[27] S. Kalyaanamoorthy, B.Q. Minh, T.K.F. Wong, A. von Haeseler, L.S. Jermiin, ModelFinder: fast model selection for accurate phylogenetic estimates, Nature Methods 14(6) (2017) 587–+.

[28] Z. Dong, P. Zhao, C. Wang, Y. Zhang, J. Chen, X. Wang, Y. Lin, Q. Xia, Comparative proteomics reveal diverse functions and dynamic changes of Bombyx mori silk proteins spun from different development stages, J Proteome Res 12(11) (2013) 5213–22.

[29] J. Wang, S. Li, Y. Zhang, H. Zheng, Z. Xu, J. Ye, J. Yu, G.K. Wong, Vertebrate gene predictions and the problem of large genes, Nat Rev Genet 4(9) (2003) 741–9.

[30] M. Levin, F. Butter, Proteotranscriptomics - A facilitator in omics research, Comput Struct Biotec 20 (2022) 3667–3675.

[31] T.T. Gai, X.L. Tong, M.J. Han, C.L. Li, C.Y. Fang, Y.L. Zou, H. Hu, H. Xiang, Z.H. Xiang, C. Lu, F.Y. Dai, Cocoonase is indispensable for Lepidoptera insects breaking the sealed cocoon, Plos Genetics 16(9) (2020).

[32] Y.R. Li, Y.K. Wei, G.Z. Zhang, Y.S. Zhang, Sericin from Fibroin-Deficient Silkworms Served as a Promising Resource for Biomedicine, Polymers-Basel 15(13) (2023).

[33] S. Prasong, S. Yaowalak, S. Wilaiwan, Characteristics of silk fiber with and without sericin component: A comparison between Bombyx mori and Philosamia ricini silks, Pak. J. Biol. Sci. 12 (2009) 872−876.

[34] Y.A. Muller, C. Heiring, R. Misselwitz, K. Welfle, H. Welfle, The cystine knot promotes folding and not thermodynamic stability in vascular endothelial growth factor, J Biol Chem 277(45) (2002) 43410–6.

[35] S.M. Hong, S.K. Nho, N.S. Kim, J.S. Lee, S.W. Kang, Gene expression profiling in the silkworm, Bombyx mori, during early embryonic development, Zoolog Sci 23(6) (2006) 517–28.

