## Supplementary material for "Unravelling the complexity of silk sericins: *P150/sericin 6* is a new silk gene in *Bombyx mori*": Table 1

| **Table 1.** List of proteins detected in *B. mori* cocoon silk by MaxQuant. Intensity is the sum of intensity values of all replicates. | | | | |  |  |  |  |
| --- | --- | --- | --- | --- | --- | --- | --- | --- |
| **Intensity** | **Accession** | **Gene** | **Residue** | **M. w. (kDa)** | **GRAVY** | **pI** | **Top3 AA** | **Signal peptide** |
| 380194000000 | XP_037869538.1 | sericin 1 isoform X1 | 3385 | 331.903 | -1.118 | 4.773 | Ser (38.405%); Gly (12.644%); Thr (9.365%) | MRFVLCCTLIALAALSVKAFG |
| 41472000000 | XP_004926112.1 | uncharacterized protein LOC101740726 | 239 | 25.909 | -0.016 | 8.824 | Leu (15.900%); Ala (10.460%); Gly (8.368%) | MKLLVVLAVCAVAAA |
| 27399600000 | XP_004922636.1 | fibrohexamerin-like | 241 | 27.201 | -0.058 | 4.555 | Leu (9.959%); Asn (7.884%); Ser (7.884%) | MKLMLCLCLLSFFGLTVA |
| 22181230000 | XP_037874919.1 | LOW QUALITY PROTEIN: zonadhesin | 3973 | 432.78 | -0.563 | 4.644 | Cys (14.372%); Gly (8.759%); Asn (8.558%) | MLKTFLFVACIANVLLVDSTEA |
| 17189800000 | NP_001108116.1 | sericin 3 precursor | 1271 | 123.298 | -1.465 | 5.708 | Ser (43.588%); Gly (11.959%); Gln (7.396%) | MNCKVALFLIVAIVAVQA |
| 11367900000 | NP_001106733.1 | fibroin heavy chain precursor | 5263 | 391.593 | 0.216 | 4.128 | Gly (45.886%); Ala (30.268%); Ser (12.065%) | MRVKTFVILCCALQYVAYTNA |
| 7049000000 | NP_001037488.1 | fibroin light chain precursor | 262 | 27.754 | 0.024 | 5.188 | Ala (13.740%); Ser (9.542%); Gly (8.397%) | MKPIFLVLLVATSAYA |
| 6734500000 | XP_004921814.1 | uncharacterized protein LOC101741789 | 474 | 50.925 | -0.448 | 5.343 | Lys (9.705%); Ser (9.494%); Val (8.228%) | MKVAAALFTILQIVNA |
| 4998900000 | NP_001037045.1 | seroin 1 precursor | 108 | 11.882 | -0.388 | 4.059 | Ser (12.037%); Asp (10.185%); Glu (9.259%) | MAFTKFLFVITLITIASA |
| 3343110000 | NP_001078833.1 | beta-N-acetylglucosaminidase 1 precursor | 611 | 69.655 | -0.287 | 6.124 | Leu (9.493%); Val (7.529%); Arg (6.710%) | MALHRLLALMTTMHLAGVLC |
| 2228430000 | XP_004921708.1 | autotransporter adhesin BpaC-like | 261 | 24.119 | -0.133 | 3.879 | Ala (29.502%); Ser (19.923%); Thr (17.241%) | MSLSRVAVFASILVCYYVSIVA |
| 1978960000 | XP_004926642.2 | apolipophorins isoform X1 | 3317 | 369.106 | -0.315 | 7.973 | Leu (9.828%); Lys (9.014%); Ser (7.266%) | MGTISFSLSVILLISVLWEPALA |
| 1776180000 | NP_001106745.1 | carboxypeptidase inhibitor precursor | 113 | 12.561 | -0.296 | 4.244 | Cys (13.274%); Glu (10.619%); Gly (8.850%) | MKFLLFCAMCILVYG |
| 1700100000 | XP_004925684.1 | rh5-interacting protein isoform X1 | 394 | 43.157 | -0.34 | 4.366 | Cys (13.959%); Pro (10.660%); Asn (8.376%) | MFILFCFISTVAA |
| 1198871000 | XP_012546338.1 | glycine-rich cell wall structural protein 1.0-like | 208 | 18.431 | -0.801 | 3.207 | Gly (36.538%); Asp (18.750%); Ala (12.500%) | MKLKWLLLLVVVAVTYVEA |
| 1102090000 | XP_037874647.1 | zonadhesin isoform X1 | 683 | 75.835 | -0.606 | 4.663 | Cys (14.641%); Glu (9.956%); Pro (7.174%) | MGRFCFIVLTAAWLVVAKADA |
| 1100470000 | XP_037867528.1 | zonadhesin isoform X1 | 2721 | 296.36 | -0.549 | 5.952 | Cys (14.627%); Pro (13.120%); Asn (8.012%) | MLVLFVGLLMLISSSTA |
| 1081719000 | XP_004928913.2 | vanin-like protein 1 | 515 | 57.684 | -0.124 | 5.014 | Val (8.932%); Leu (7.573%); Ser (7.379%) | MKLIFYGLLIFSLTKATS |
| 1060700000 | XP_004923912.1 | alcohol dehydrogenase 2 | 319 | 35.344 | -0.202 | 6.788 | Val (9.404%); Ile (8.464%); Asn (8.464%) | MFVITYFNILFVALFFVNVTST |
| 1034207000 | XP_021204310.2 | uncharacterized protein LOC101744556 isoform X1 | 1979 | 222.161 | -0.727 | 4.753 | Asn (9.803%); Pro (8.489%); Ser (8.439%) | MEYKVNLIFLVVFSSKIVVSIP |
| 961800000 | XP_037871220.1 | fibrohexamerin-like | 240 | 27.327 | -0.179 | 4.591 | Leu (9.583%); Asn (8.333%); Glu (7.500%) | MRLVWYSCLALFGLITA |
| 880960000 | XP_037873722.1 | glucose dehydrogenase [FAD, quinone] | 577 | 64.896 | -0.155 | 7.01 | Leu (10.052%); Val (8.146%); Lys (7.452%) | - |
| 749953000 | XP_037869877.1 | uncharacterized protein LOC101740721 isoform X1 | 2270 | 251.441 | -0.988 | 6.716 | Lys (9.824%); Thr (9.604%); Ser (9.471%) | MDWTKYLVLFILMVVVSS |
| 717188000 | XP_037874060.1 | myrosinase 1 | 567 | 65.135 | -0.404 | 6.173 | Leu (8.466%); Ser (8.113%); Ala (7.760%) | MFGLLSMWISWDLFLFERPASV |
| 700756000 | NP_001139413.1 | fibrohexamerin precursor | 220 | 25.167 | -0.102 | 7.162 | Leu (10.000%); Ala (7.273%); Phe (6.818%) | MLARCLAVAAVAVLASA |
| 677915000 | NP_001108339.1 | integument esterase 1 precursor | 561 | 63.237 | -0.425 | 6.835 | Leu (8.021%); Gly (7.843%); Pro (7.843%) | MLLKLIFICAIVYYADA |
| 673315000 | XP_012551240.1 | uncharacterized protein LOC105842476 | 234 | 26.439 | -0.007 | 4.168 | Leu (9.402%); Asn (8.547%); Val (7.692%) | MEFELCLCILSVLGLIAA |
| 474186000 | XP_004924748.1 | proton-coupled folate transporter | 508 | 57.334 | 0.229 | 8.584 | Leu (10.827%); Ile (9.646%); Ser (8.661%) | - |
| 456522000 | XP_012546430.1 | lysosomal acid glucosylceramidase isoform X1 | 536 | 61.151 | -0.243 | 7.051 | Leu (9.142%); Lys (7.836%); Ile (7.090%) | - |
| 403273500 | XP_004928916.1 | glucose dehydrogenase [FAD, quinone] | 657 | 73.389 | -0.294 | 8.726 | Leu (8.676%); Val (7.610%); Gly (7.154%) | MHTRRKIIFMWFLLFVNANCCDA |
| 352649000 | XP_037868215.1 | beta-glucuronidase isoform X1 | 688 | 79.06 | -0.304 | 7.015 | Val (7.849%); Leu (6.977%); Thr (6.831%) | - |
| 342382000 | XP_004927001.2 | lipase 1 | 399 | 46.491 | -0.293 | 5.885 | Leu (9.023%); Glu (8.020%); Ile (7.519%) | MVKIGLRTFFMILSIGLHLASA |
| 342254000 | XP_004922749.1 | protein D2 | 209 | 23.055 | 0.077 | 6.528 | Val (11.005%); Leu (10.048%); Ala (8.612%) | MGALIRCAIVLLTVATIVDFRVLTRA |
| 319436000 | XP_037872749.1 | dehydrogenase mpl7 isoform X1 | 585 | 65.476 | -0.206 | 9.092 | Lys (9.573%); Leu (9.060%); Val (8.547%) | - |
| 314065000 | NP_001037046.2 | seroin 2 precursor | 112 | 12.273 | -0.287 | 9.883 | Ser (12.500%); Lys (10.714%); Pro (10.714%) | MAFTKFLFMLSLITIASA |
| 298483000 | XP_037869036.1 | beta-galactosidase isoform X1 | 2942 | 328.542 | -0.211 | 6.857 | Leu (9.007%); Val (7.988%); Gly (7.104%) | MTLLIATFALLCFLGQTYS |
| 283373000 | XP_012545566.1 | yellow-d isoform X1 | 475 | 53.869 | -0.208 | 6.511 | Val (8.211%); Leu (7.158%); Ser (7.158%) | - |
| 252201950 | NP_001036850.1 | cathepsin B precursor | 337 | 37.556 | -0.441 | 6.382 | Gly (9.792%); Ser (7.122%); Asp (6.825%) | MFISRAAYVTLVCVLAAA |
| 245267000 | XP_004924612.2 | esterase FE4 | 550 | 61.673 | -0.231 | 5.06 | Leu (8.000%); Gly (7.636%); Ile (7.273%) | MLIKTFFVIASVVYVLG |
| 181921000 | XP_037876844.1 | latent-transforming growth factor beta-binding protein 4 | 1359 | 150.321 | -0.652 | 5.589 | Thr (10.302%); Ser (8.389%); Cys (7.873%) | MKFIVVIFTFLFSTVNA |
| 165783000 | XP_004929234.1 | venom dipeptidyl peptidase 4 | 740 | 83.919 | -0.199 | 5.527 | Leu (8.243%); Val (7.973%); Ile (7.162%) | MAMEQAVFLLMGLICHTQSSA |
| 149592000 | NP_001093080.1 | ecdysteroid-regulated 16 kDa protein precursor | 145 | 15.824 | 0.179 | 6.336 | Leu (11.724%); Val (8.276%); Ala (7.586%) | MLFFITAAVLLASAEA |
| 148759000 | XP_037874096.1 | myrosinase 1 isoform X1 | 495 | 57.471 | -0.473 | 5.019 | Leu (8.081%); Glu (7.879%); Ser (7.071%) | MNLSWQVAFSALMACAWG |
| 145757000 | XP_037868236.1 | juvenile hormone esterase-like | 562 | 63.516 | -0.189 | 9.306 | Leu (9.253%); Asn (8.541%); Ile (7.295%) | MSNTFYLITVLILVLNTINA |
| 143543000 | NP_001268822.1 | neutral alpha-glucosidase AB-like precursor | 925 | 104.635 | -0.371 | 5.706 | Leu (8.432%); Val (8.324%); Ala (8.000%) | MKVWALVLVAAFVIIGISA |
| 137424000 | XP_004926880.2 | lysosomal acid glucosylceramidase isoform X1 | 530 | 59.677 | -0.19 | 4.426 | Leu (8.113%); Asp (7.358%); Ile (7.170%) | MDINKCIISASLWAAVYLLFGSADA |
| 134477000 | XP_012547740.1 | heat shock 70 kD protein cognate isoform X1 | 960 | 106.18 | -0.4 | 5.486 | Lys (8.542%); Val (8.229%); Ala (8.125%) | - |
| 121943000 | XP_037868883.1 | arylsulfatase B | 537 | 60.465 | -0.397 | 6.796 | Leu (8.752%); Gly (8.194%); Lys (7.635%) | MFVFRVLLISVLMLVAES |
| 116001000 | XP_012544831.3 | uncharacterized protein LOC101741510 | 1999 | 228.101 | -0.53 | 5.916 | Val (8.054%); Leu (8.004%); Lys (7.404%) | MSIKSAYLLIFQIIVIYNCSC |
| 114229100 | NP_001037047.1 | silk proteinase inhibitor precursor | 65 | 6.962 | 0.554 | 8.134 | Cys (13.846%); Gly (10.769%); Thr (10.769%) | MKTSIVLIFLLVACCTLGAES |
| 113920600 | XP_021206228.1 | poly(U)-specific endoribonuclease-D isoform X1 | 536 | 60.005 | -0.79 | 9.886 | Ser (10.634%); Thr (9.888%); Asn (8.769%) | - |
| 102791000 | NP_001106744.1 | antennal binding protein precursor | 140 | 15.49 | -0.11 | 7.206 | Lys (11.429%); Ala (10.000%); Leu (7.857%) | MMYLSFVVLICLAFAVFNCGA |
| 97291000 | XP_004927018.1 | C3 and PZP-like alpha-2-macroglobulin domain-containing protein 8 | 1448 | 155.934 | -0.031 | 5.938 | Ala (11.119%); Leu (10.152%); Ser (8.840%) | MIEKIKTLLLFILSVPAVTQC |
| 96514000 | XP_037874655.1 | inducible metalloproteinase inhibitor protein-like isoform X1 | 200 | 22.342 | -0.704 | 5.769 | Cys (10.500%); Glu (9.500%); Ala (7.000%) | - |
| 93201000 | XP_021203913.1 | uncharacterized protein LOC101747119 isoform X1 | 722 | 80.01 | -0.235 | 6.67 | Val (9.695%); Leu (7.895%); Ser (7.895%) | MNSNEIIYFVILTLCSSVAG |
| 77323000 | XP_012545193.1 | uncharacterized protein LOC101735738 isoform X1 | 671 | 75.143 | -0.309 | 6.349 | Leu (10.581%); Gly (8.197%); Val (7.899%) | MSVRFVRTILVLLAIVHNSRA |
| 75014000 | NP_001040174.1 | alpha-esterase 13 precursor | 540 | 61.416 | -0.241 | 5.012 | Leu (9.815%); Phe (7.593%); Lys (7.593%) | MLFAIIICVQVLSVFG |
| 74413000 | XP_037876352.1 | 4-hydroxy-tetrahydrodipicolinate synthase-like | 328 | 35.694 | 0.094 | 4.9 | Leu (11.585%); Ala (8.537%); Ile (7.927%) | MYSISGLTFLCVLIINVFDTKC |
| 73489010 | XP_037867171.1 | 15-hydroxyprostaglandin dehydrogenase [NAD(+)]-like | 289 | 31.605 | -0.025 | 9.45 | Lys (10.727%); Ile (10.035%); Ala (7.958%) | MTLSLLKTFFVSCFIINIVSA |
| 71931700 | NP_001037075.1 | calreticulin precursor | 398 | 45.802 | -0.992 | 4.245 | Asp (13.819%); Lys (12.060%); Glu (11.307%) | MKAVVLVVVSLLALSSINC |
| 68856900 | XP_021206073.2 | uncharacterized protein LOC105842244 | 1131 | 127.974 | -0.974 | 9.852 | Thr (10.698%); Ser (9.991%); Pro (9.107%) | MCGARLLLAATALQALILAVSC |
| 67160000 | XP_012547984.1 | uncharacterized protein LOC101744060 | 934 | 101.918 | -0.275 | 4.433 | Leu (9.850%); Ser (8.565%); Ala (8.458%) | MNANKKAIVLLTLINILLVRS |
| 66627000 | XP_037874369.1 | fibrillin-1 isoform X1 | 3047 | 325.331 | -0.468 | 4.496 | Gly (12.504%); Cys (12.143%); Asp (7.844%) | MGDGAKMSYWRLLVAAALLASVAG |
| 60053000 | XP_012550868.1 | uncharacterized protein LOC778506 isoform X1 | 292 | 30.66 | -0.599 | 4.054 | Ala (16.781%); Glu (12.329%); Lys (11.644%) | - |
| 59817000 | XP_037868167.1 | uncharacterized protein LOC101736658 isoform X1 | 456 | 51.569 | -0.228 | 6.318 | Leu (9.649%); Ser (8.333%); Val (8.114%) | MGSIETLVLLQLVYISSC |
| 59417900 | XP_004929039.1 | phospholipase A2 | 177 | 20.411 | -0.24 | 4.652 | Glu (7.910%); Asp (7.345%); Gly (7.345%) | MFRHLFCLIVLIAVHTNKA |
| 58891600 | XP_037867041.1 | alpha-crystallin B chain | 241 | 26.543 | -0.263 | 4.361 | Val (11.618%); Ala (9.129%); Thr (9.129%) | MFSPRLFAVFIAFAGFATVTA |
| 55656900 | XP_021207905.1 | uncharacterized protein LOC101743953 isoform X1 | 472 | 52.872 | -0.063 | 7.143 | Leu (11.441%); Ile (8.263%); Ala (8.051%) | MAVSLYVVLVLATTALG |
| 55294500 | XP_037868227.1 | mucin-5AC | 1768 | 198.075 | -0.847 | 6.971 | Thr (14.989%); Ser (8.484%); Glu (8.258%) | MGIFDYKFIGKLFKMESTIWLTLLLAVIATTPVSS |
| 50392000 | NP_001037090.1 | serine protease inhibitor 4 precursor | 410 | 46.3 | -0.095 | 7.322 | Leu (9.024%); Ser (7.805%); Ile (7.561%) | MCLLKFLVLAIVPLSFA |
| 44188900 | NP_001037171.1 | protein disulfide isomerase precursor | 494 | 55.589 | -0.278 | 4.325 | Glu (11.134%); Ala (8.704%); Lys (8.502%) | MRVLIFTAIALLGLALG |
| 41498000 | NP_001119727.1 | actin, cytoplasmic A4 | 376 | 41.822 | -0.193 | 5.156 | Ala (7.979%); Glu (7.447%); Gly (7.447%) | - |
| 40515000 | XP_037874307.1 | basement membrane-specific heparan sulfate proteoglycan core protein isoform X1 | 4228 | 463.254 | -0.538 | 4.432 | Gly (8.609%); Ser (8.538%); Asp (7.427%) | MRAGGLLAAALLLLSTFTIQVLKA |
| 39652000 | XP_004926666.2 | alaserpin | 410 | 45.953 | -0.194 | 5.018 | Leu (10.244%); Ile (9.024%); Glu (8.293%) | - |
| 38903380 | XP_012551293.2 | heme peroxidase 2 | 1344 | 149.897 | -0.304 | 6.792 | Leu (10.045%); Ala (8.929%); Pro (7.068%) | MGAFKASIRSLLLFTLALT |
| 37924900 | NP_001155191.2 | silk gland derived serine protease 1 precursor | 392 | 42.098 | 0.134 | 7.393 | Gly (9.694%); Ser (9.439%); Val (8.163%) | MCLQLVLVVLALNGVLS |
| 37469400 | NP_001040479.1 | peptidylprolyl isomerase B precursor | 205 | 22.397 | -0.267 | 8.842 | Gly (12.195%); Lys (10.244%); Thr (8.780%) | MGTLTMALGILLFIASAKS |
| 37262200 | NP_001037430.1 | yellow-b precursor | 457 | 50.803 | -0.156 | 5.119 | Leu (9.409%); Ala (8.972%); Asn (7.440%) | MRYTLTSLSMMIAIVLLAAVAAAAA |
| 36225300 | NP_001037073.1 | glucosidase precursor | 491 | 55.596 | -0.375 | 4.575 | Asp (7.943%); Leu (7.943%); Gly (7.536%) | MAWLTTLSILAVCHTGLA |
| 34377200 | NP_001268827.1 | low molecular mass 30 kDa lipoprotein 21G1-like precursor | 256 | 29.858 | -0.714 | 4.942 | Asp (10.938%); Lys (8.984%); Ser (8.594%) | MKLKILTTLSVIIATFFNMSVEA |
| 30436100 | XP_037870111.1 | probable polyamine oxidase 5 isoform X1 | 485 | 53.978 | -0.216 | 6.632 | Leu (9.485%); Gly (7.835%); Thr (7.835%) | MIVRVAILVLCVAVQGRG |
| 29882500 | NP_001036944.1 | juvenile hormone binding protein an-0895 precursor | 243 | 27.294 | -0.047 | 4.558 | Leu (13.992%); Val (9.053%); Asn (8.642%) | MWTGLFLVLGLYESVVG |
| 27055300 | XP_021205588.1 | probable cyclin-dependent serine/threonine-protein kinase DDB_G0292550 isoform X2 | 544 | 59.257 | -1.054 | 6.833 | Ser (18.199%); Asn (14.154%); Asp (8.272%) | MNIARSIIFSVYVILLVGLSPTSG |
| 25743100 | XP_004926879.1 | apolipoprotein D | 269 | 29.851 | -0.199 | 6.508 | Val (8.922%); Ser (8.550%); Leu (8.178%) | MWKLTVLGVFLTVTYVYS |
| 25471600 | XP_004926990.1 | ATP-dependent (S)-NAD(P)H-hydrate dehydratase | 327 | 35.913 | -0.099 | 7.438 | Ser (10.092%); Ile (9.786%); Gly (8.257%) | MTPSFNIILVILQVILFTFQITNG |
| 25363500 | XP_037876745.1 | calsyntenin-1 isoform X1 | 953 | 105.096 | -0.324 | 5.94 | Ala (9.654%); Val (8.919%); Leu (8.185%) | MIVRFLSVLCVGVFLTAVYG |
| 23648900 | XP_004923234.1 | putative alpha-L-fucosidase | 477 | 55.387 | -0.509 | 7.75 | Leu (7.757%); Ser (7.128%); Gly (6.918%) | MKSFFLVFLFGFVNG |
| 20809000 | XP_004933848.1 | 15-hydroxyprostaglandin dehydrogenase [NAD(+)] | 283 | 31.156 | -0.108 | 6.252 | Ile (8.834%); Ala (8.481%); Lys (7.774%) | MAVELTFLFCLVILSVRA |
| 19455700 | NP_001091826.1 | ubiquitin and ribosomal protein S27a | 155 | 17.847 | -0.797 | 10.483 | Lys (16.129%); Leu (7.742%); Gly (7.097%) | - |
| 19071700 | NP_001166287.1 | sericin 2 isoform 1 precursor | 1758 | 198.657 | -2.152 | 9.158 | Lys (17.179%); Ser (15.131%); Asp (11.832%) | MKIPYVLLFLVGVAVVNA |
| 17735800 | XP_004928346.2 | uncharacterized protein LOC101735388 | 111 | 12.342 | -0.008 | 5.119 | Ile (9.910%); Asp (9.009%); Cys (8.108%) | MFSRLVVLGFVICVANA |
| 17105900 | XP_037867498.1 | chaoptin isoform X1 | 1367 | 155.077 | -0.135 | 6.335 | Leu (15.435%); Ser (8.925%); Asn (7.901%) | MSLMAVVKFGYTLVAVTLLLMIWASLSRA |
| 16636200 | P150 | P150 | 4552 | 467.241 | -0.672 | 3.815 | Thr (27.307%); Ser (14.455%); Ala (12.017%) | MKVLCAIVLYIALMQPALC |
| 16537500 | XP_004921556.2 | mucin-5AC isoform X1 | 856 | 92.849 | -0.557 | 4.199 | Glu (10.748%); Ser (10.748%); Thr (10.397%) | MIRSKGDMASPRFSTILLAAFHLFTVCWG |
| 16068900 | XP_004923220.1 | endoplasmic reticulum resident protein 44 | 410 | 47.029 | -0.391 | 6.032 | Leu (10.000%); Lys (8.537%); Glu (7.561%) | MLKMNWKCPNLFNFKLSSCLFILLCHSFYNPTDS |
| 14948100 | XP_037872311.1 | nardilysin | 1146 | 132.148 | -0.49 | 6.333 | Leu (9.686%); Glu (8.551%); Ser (8.115%) | - |
| 14014160 | XP_004932517.1 | extracellular serine/threonine protein CG31145 isoform X1 | 579 | 66.067 | -0.285 | 7.713 | Leu (10.363%); Arg (8.117%); Ala (7.772%) | - |
| 12870000 | NP_001243928.1 | heat shock protein 68 | 628 | 69.367 | -0.436 | 5.674 | Ala (9.713%); Asp (7.643%); Leu (7.643%) | - |
| 12693000 | XP_037869639.1 | alpha-1,6-mannosyl-glycoprotein 2-beta-N-acetylglucosaminyltransferase isoform X1 | 522 | 60.708 | -0.538 | 8.814 | Leu (8.238%); Asn (7.088%); Val (6.513%) | - |
| 12590400 | XP_021205679.2 | mucin-12 isoform X1 | 3173 | 368.381 | -1.498 | 4.137 | Thr (18.374%); Glu (16.451%); Gln (15.065%) | MTAKTCCLLWVIVLASILATVVT |
| 11544300 | XP_004930613.1 | peroxidase isoform X1 | 799 | 90.376 | -0.4 | 6.855 | Leu (9.387%); Ser (8.385%); Thr (6.383%) | - |
| 10500500 | XP_004930766.1 | lachesin isoform X1 | 370 | 41.237 | -0.2 | 7.444 | Ile (8.378%); Ala (8.108%); Val (7.297%) | MDRNIYVYVFYTLLLYFTCNVSA |
| 10166180 | XP_012544866.1 | zinc finger CCCH domain-containing protein 13 | 1004 | 117.836 | -1.437 | 8.884 | Arg (13.247%); Pro (11.554%); Thr (9.661%) | MKRSRHRWLGFTIFVCLLAILASELPNTAA |
| 9468900 | XP_004925158.1 | mannosyl-oligosaccharide glucosidase | 807 | 91.5 | -0.417 | 9.002 | Leu (10.905%); Gly (8.302%); Ser (6.939%) | - |
| 9318700 | XP_037871889.1 | chitinase-like protein EN03 isoform X1 | 455 | 50.569 | -0.292 | 7.002 | Leu (11.209%); Ala (8.132%); Gly (7.692%) | MKLFIALVGLLALAKA |
| 8995100 | NP_001037584.1 | ferritin precursor | 229 | 26.033 | -0.359 | 7.28 | Leu (11.790%); Ala (7.860%); Lys (7.860%) | MKVYALIVACLALGVLA |
| 8964600 | XP_037869963.1 | low-density lipoprotein receptor-related protein 2 isoform X1 | 4625 | 512.561 | -0.477 | 5.013 | Asp (9.319%); Gly (7.892%); Cys (6.962%) | MRSAREAFCLLLLAAAAAA |
| 7899300 | XP_004925541.1 | uncharacterized protein LOC100500745 | 110 | 12.313 | 0.187 | 9.144 | Ile (10.909%); Leu (10.909%); Lys (8.182%) | MKFAPVLIFIFLSLVCVIQC |
| 7344340 | NP_001037486.1 | low molecular 30 kDa lipoprotein PBMHP-6 precursor | 256 | 29.734 | -0.444 | 6.523 | Lys (9.375%); Leu (7.812%); Glu (7.422%) | MRLTLFAFVLAVCALASNA |
| 6479200 | NP_001153666.1 | histone H2A-like protein 2 | 124 | 13.377 | -0.318 | 11.361 | Ala (12.097%); Leu (12.097%); Gly (11.290%) | - |
| 6435200 | NP_001243674.1 | thrombospondin type-1 domain-containing protein 4-like precursor | 574 | 62.763 | -0.464 | 8.876 | Arg (9.059%); Gly (8.885%); Ser (8.885%) | MAGADLYLLFVIFVTMVVG |
| 5897400 | XP_004929845.2 | peroxidasin isoform X1 | 1379 | 157.181 | -0.35 | 5.959 | Leu (10.007%); Asp (6.744%); Glu (6.526%) | MRLCNFFKFLLIFNLFSLILS |
| 4868770 | XP_037871221.1 | fibrohexamerin-like | 235 | 26.815 | -0.281 | 4.483 | Leu (10.638%); Ser (7.660%); Asp (7.234%) | MKYKLCLFLSSLLNLIAA |
| 4099550 | NP_001037351.1 | cathepsin D precursor | 384 | 41.623 | 0.154 | 6.732 | Leu (9.896%); Ala (9.375%); Gly (9.115%) | MGKISLFFLALIASSVMA |
| 3173900 | XP_012549547.1 | malate dehydrogenase isoform X1 | 645 | 70.89 | -0.186 | 6.641 | Leu (9.457%); Ala (8.992%); Gly (8.992%) | - |
| 3103050 | XP_012551451.2 | uncharacterized protein LOC105842505 | 1097 | 122.723 | -0.805 | 6.678 | Pro (12.124%); Thr (8.204%); Ser (7.840%) | MVRGATLALTLALAWVCVVRT |
| 2356030 | NP_001037057.1 | BCP inhibitor precursor | 105 | 12.249 | -0.735 | 6.519 | Glu (9.524%); Leu (9.524%); Ala (8.571%) | MNFVSVALLIATVVMASSA |
| 1717200 | XP_004925279.1 | uncharacterized protein LOC101741292 | 235 | 27.212 | -0.601 | 8.743 | Arg (10.213%); Leu (9.787%); Ser (8.085%) | MGARGCISCLLLLTCGQIVRS |
