## Supplements for "Unravelling the complexity of silk sericins: *P150/sericin 6* is a new silk gene in *Bombyx mori*"

**Supplementary Figure S1.** Expression levels of *B. mori* P150 gene along with major silk genes (Sericin 1-3 and Mucin-12) and the control gene (mothers against dpp; Mad) in different parts of silk glands. Heat maps represent the log2 fold-change values. As shown in both figures, P150 is highly expressed in middle silk glands. Sources of the raw RNA-seq data were listed in **Supplementary Table S1**.

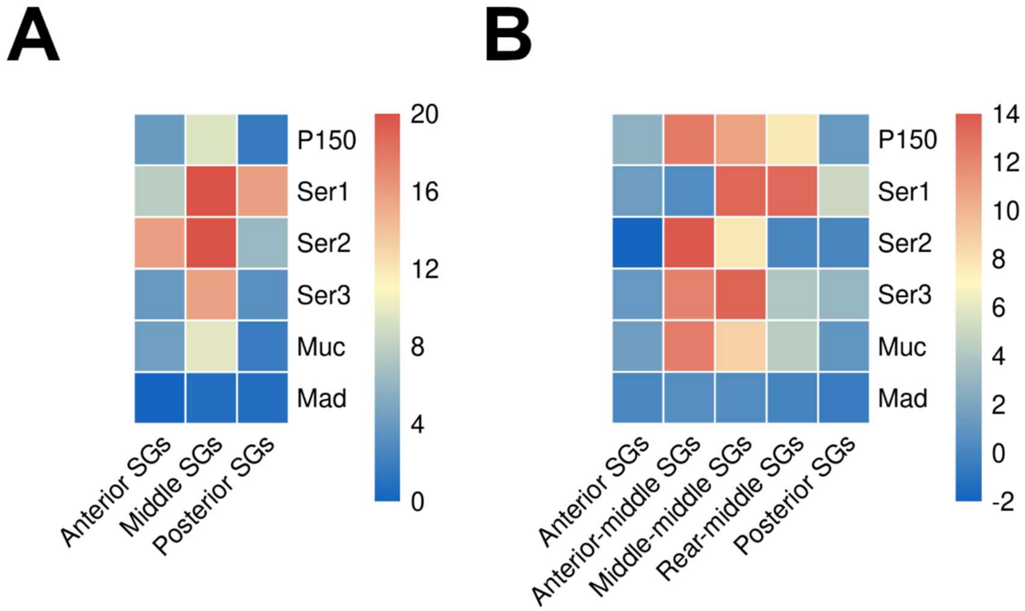

**Supplementary Table S1.** Sources of the RNA-seq datasets utilized to verify the expression of *B. mori* P150 and other genes.

(A) datasets used in **Supplementary Figure S1A**

| Run | BioProject | BioSample | Age | Instrument | LibraryLayout | LibrarySource | Organism | Tissue |
|---|---|---|---|---|---|---|---|---|
| SRR10035619 | PRJNA559726 | SAMN12608940 | 5th instar day 3 | Illumina NovaSeq 6000 | PAIRED | TRANSCRIPTOMIC | Bombyx mori | Posterior-Silk-Gland |
| SRR10035620 | PRJNA559726 | SAMN12608939 | 5th instar day 3 | Illumina NovaSeq 6000 | PAIRED | TRANSCRIPTOMIC | Bombyx mori | Posterior-Silk-Gland |
| SRR10035621 | PRJNA559726 | SAMN12608938 | 5th instar day 3 | Illumina NovaSeq 6000 | PAIRED | TRANSCRIPTOMIC | Bombyx mori | Posterior-Silk-Gland |
| SRR10035669 | PRJNA559726 | SAMN12609122 | 5th instar day 3 | Illumina NovaSeq 6000 | PAIRED | TRANSCRIPTOMIC | Bombyx mori | Middle-Silk-Gland |
| SRR10035670 | PRJNA559726 | SAMN12609121 | 5th instar day 3 | Illumina NovaSeq 6000 | PAIRED | TRANSCRIPTOMIC | Bombyx mori | Middle-Silk-Gland |
| SRR10035671 | PRJNA559726 | SAMN12609120 | 5th instar day 3 | Illumina NovaSeq 6000 | PAIRED | TRANSCRIPTOMIC | Bombyx mori | Middle-Silk-Gland |
| SRR10035726 | PRJNA559726 | SAMN12609071 | 5th instar day 3 | Illumina NovaSeq 6000 | PAIRED | TRANSCRIPTOMIC | Bombyx mori | Midgut |
| SRR10035727 | PRJNA559726 | SAMN12609070 | 5th instar day 3 | Illumina NovaSeq 6000 | PAIRED | TRANSCRIPTOMIC | Bombyx mori | Midgut |
| SRR10035728 | PRJNA559726 | SAMN12609069 | 5th instar day 3 | Illumina NovaSeq 6000 | PAIRED | TRANSCRIPTOMIC | Bombyx mori | Midgut |
| SRR10035756 | PRJNA559726 | SAMN12609044 | 5th instar day 3 | Illumina NovaSeq 6000 | PAIRED | TRANSCRIPTOMIC | Bombyx mori | Anterior-Silk-Gland |
| SRR10035757 | PRJNA559726 | SAMN12609043 | 5th instar day 3 | Illumina NovaSeq 6000 | PAIRED | TRANSCRIPTOMIC | Bombyx mori | Anterior-Silk-Gland |
| SRR10035758 | PRJNA559726 | SAMN12609042 | 5th instar day 3 | Illumina NovaSeq 6000 | PAIRED | TRANSCRIPTOMIC | Bombyx mori | Anterior-Silk-Gland |

(B) datasets used in **Supplementary Figure S1B**

| Run | BioProject | BioSample | Age | Instrument | LibraryLayout | LibrarySource | Organism | Tissue |
|---|---|---|---|---|---|---|---|---|
| DRR186474 | PRJDB8614 | SAMD00180408 | fifth instar day 3 | Illumina NovaSeq 6000 | PAIRED | TRANSCRIPTOMIC | Bombyx mori | Anterior silk gland |
| DRR186475 | PRJDB8614 | SAMD00180409 | fifth instar day 3 | Illumina NovaSeq 6000 | PAIRED | TRANSCRIPTOMIC | Bombyx mori | Anterior silk gland |
| DRR186476 | PRJDB8614 | SAMD00180410 | fifth instar day 3 | Illumina NovaSeq 6000 | PAIRED | TRANSCRIPTOMIC | Bombyx mori | Anterior silk gland |
| DRR186477 | PRJDB8614 | SAMD00180411 | fifth instar day 3 | Illumina NovaSeq 6000 | PAIRED | TRANSCRIPTOMIC | Bombyx mori | Anterior part of the middle silk gland |
| DRR186478 | PRJDB8614 | SAMD00180412 | fifth instar day 3 | Illumina NovaSeq 6000 | PAIRED | TRANSCRIPTOMIC | Bombyx mori | Anterior part of the middle silk gland |
| DRR186479 | PRJDB8614 | SAMD00180413 | fifth instar day 3 | Illumina NovaSeq 6000 | PAIRED | TRANSCRIPTOMIC | Bombyx mori | Anterior part of the middle silk gland |
| DRR186480 | PRJDB8614 | SAMD00180414 | fifth instar day 3 | Illumina NovaSeq 6000 | PAIRED | TRANSCRIPTOMIC | Bombyx mori | Middle part of the middle silk gland |
| DRR186481 | PRJDB8614 | SAMD00180415 | fifth instar day 3 | Illumina NovaSeq 6000 | PAIRED | TRANSCRIPTOMIC | Bombyx mori | Middle part of the middle silk gland |
| DRR186482 | PRJDB8614 | SAMD00180416 | fifth instar day 3 | Illumina NovaSeq 6000 | PAIRED | TRANSCRIPTOMIC | Bombyx mori | Middle part of the middle silk gland |
| DRR186483 | PRJDB8614 | SAMD00180417 | fifth instar day 3 | Illumina NovaSeq 6000 | PAIRED | TRANSCRIPTOMIC | Bombyx mori | Posterior part of the middle silk gland |
| DRR186484 | PRJDB8614 | SAMD00180418 | fifth instar day 3 | Illumina NovaSeq 6000 | PAIRED | TRANSCRIPTOMIC | Bombyx mori | Posterior part of the middle silk gland |
| DRR186485 | PRJDB8614 | SAMD00180419 | fifth instar day 3 | Illumina NovaSeq 6000 | PAIRED | TRANSCRIPTOMIC | Bombyx mori | Posterior part of the middle silk gland |
| DRR186486 | PRJDB8614 | SAMD00180420 | fifth instar day 3 | Illumina NovaSeq 6000 | PAIRED | TRANSCRIPTOMIC | Bombyx mori | Posterior silk gland |
| DRR186487 | PRJDB8614 | SAMD00180421 | fifth instar day 3 | Illumina NovaSeq 6000 | PAIRED | TRANSCRIPTOMIC | Bombyx mori | Posterior silk gland |
| DRR186488 | PRJDB8614 | SAMD00180422 | fifth instar day 3 | Illumina NovaSeq 6000 | PAIRED | TRANSCRIPTOMIC | Bombyx mori | Posterior silk gland |
| DRR186492 | PRJDB8614 | SAMD00180426 | fifth instar day 3 | Illumina NovaSeq 6000 | PAIRED | TRANSCRIPTOMIC | Bombyx mori | Midgut |
| DRR186493 | PRJDB8614 | SAMD00180427 | fifth instar day 3 | Illumina NovaSeq 6000 | PAIRED | TRANSCRIPTOMIC | Bombyx mori | Midgut |
| DRR186494 | PRJDB8614 | SAMD00180428 | fifth instar day 3 | Illumina NovaSeq 6000 | PAIRED | TRANSCRIPTOMIC | Bombyx mori | Midgut |

**Supplementary Table S2.** Sequences of primers used in (A) exon junction sequencing and (B) qPCR analysis.

| (A) Sequencing primers | |
|---|---|
| P150 #1F | GAATCGAGCACTGCAGGAGT |
| P150 #1R | AGTAGGCTGATCGTGGACCT |
| P150 #2F | CCACGAAACAGTTTGCTCCA |
| P150 #2R | TAGGCTCGATTGTGGACTGC |
| P150 #3F | CCCAAGGCTTCGGTGATGAT |
| P150 #3R | TATGCATTGCCGGAGCTTGA |

| (B) qPCR primers | |
|---|---|
| P150 F | CCCAAGGCTTCGGTGATGAT |
| P150 R | TATGCATTGCCGGAGCTTGA |
| Ser1 F | CACAACCGATAAGACGAG |
| Ser1 R | GACGAAGTGGAGGAAGC |
| Ser2 F | CATCGGCTGACTACCA |
| Ser2 R | AGAGTTGCTGCCCTTAC |
| Ser3 F | TGTCTCGTCGGTGGAA |
| Ser3 R | TTGTTGTATGACTGGCTCT |
| Mad F | ACACAAGGCGTCACATAGGG |
| Mad R | TGGTGATTGCAGTTACGGCT |
| EF1a F | CAAGTCTGGAGATGCAGCCA |
| EF1a R | GGGGTGGGAATTCCTGGAAG |

**Supplementary Table S3.** Statistical analysis of quantitative PCR analysis of gene expression of genes in different tissues. Statistical differences were evaluated by Student´s t-test. ASG: anterior silk glands; MSG: middle silk glands; PSG: posterior silk glands; OVA: ovaries. Backgound colors indicate statistical significance: orange ($p$ < 0.001); yellow ($p$ < 0.01); blue ($p$ < 0.05); blank ($p$ >=0.05).

| | ASG | | | | | MSG | | | | | PSG | | | | | OVA | | | | | GUT | | | | |
|---|---|---|---|---|---|---|---|---|---|---|---|---|---|---|---|---|---|---|---|---|---|---|---|---|---|
| **ASG** | | | | | | | | | | | | | | | | | | | | | | | | | |
| **MSG** | 6.89E-03 | 9.38E-05 | 5.64E-02 | 3.66E-01 | 8.29E-01 | | | | | | | | | | | | | | | | | | | | |
| **PSG** | 1.16E-01 | 3.66E-02 | 8.92E-01 | 8.28E-02 | 8.81E-01 | 1.14E-04 | 1.98E-02 | 1.07E-03 | 9.82E-02 | 8.89E-01 | | | | | | | | | | | | | | | |
| **OVA** | 3.89E-01 | 4.14E-02 | 1.84E-01 | 4.36E-01 | 1.22E-01 | 2.57E-06 | 3.05E-04 | 1.37E-07 | 2.23E-01 | 1.99E-03 | 2.64E-02 | 2.54E-02 | 7.48E-03 | 6.18E-03 | 9.12E-03 | | | | | | | | | | |
| **GUT** | 2.28E-01 | 1.06E-02 | 1.50E-01 | 1.59E-02 | 4.47E-01 | 4.10E-05 | 1.20E-04 | 3.08E-08 | 2.81E-02 | 1.96E-01 | 1.13E-01 | 1.80E-02 | 4.87E-03 | 1.10E-01 | 2.42E-01 | 1.40E-01 | 7.94E-02 | 4.12E-03 | 4.05E-02 | 2.71E-03 | | | | | |
| | **P150** | **Ser1** | **Ser2** | **Ser3** | **Muc** | **P150** | **Ser1** | **Ser2** | **Ser3** | **Muc** | **P150** | **Ser1** | **Ser2** | **Ser3** | **Muc** | **P150** | **Ser1** | **Ser2** | **Ser3** | **Muc** | **P150** | **Ser1** | **Ser2** | **Ser3** | **Muc** |

**Supplementary Table S4.** Properties of 2 types of repeat sequences in *B. mori* P150 protein. GRAVY: grand average of hydropathy; M. w.: molecular weight; pI: isoelectric point.

| | GRAVY | Residue | M. w. (Da) | pI | Top1 AA | Top2 AA | Top3 AA | Top4 AA | Top5 AA |
|---|---|---|---|---|---|---|---|---|---|
| Repeat 1 | -0.8512 | 1275 | 133254.12 | 3.8738 | Thr: 417 (32.706%) | Ser: 237 (18.588%) | Glu: 162 (12.706%) | Pro: 75 (5.882%) | Val: 75 (5.882%) |
| Repeat 2 | -0.4541 | 2220 | 215927.31 | 2.6857 | Thr: 696 (31.351%) | Ala: 440 (19.820%) | Ser: 320 (14.414%) | Glu: 211 (9.505%) | Val: 161 (7.252%) |

**Supplementary Table S5.** Properties of *B. mori* sericins and mucin-12 proteins. GRAVY: grand average of hydropathy; M. w.: molecular weight; pI: isoelectric point. The conserved cystein residues at the C-terminal region were labeled in red.

| Gene | Accession | GRAVY | Residue | M. w. (Da) | pI | Top1 AA | Top2 AA | Top3 AA | Top4 AA | Top5 AA | C-terminus |
|---|---|---|---|---|---|---|---|---|---|---|---|
| Sericin 1 | XP_037869538.1 | -1.1181 | 3385 | 331903.34 | 4.7726 | Ser: 1300 (38.405%) | Gly: 428 (12.644%) | Thr: 317 (9.365%) | Asn: 270 (7.976%) | Asp: 221 (6.529%) | LLHKPGQGKICLCFENIFDIPYHLRKNIGV |
| Sericin 2 | NP_001166287.1 | -2.1516 | 1758 | 198657.39 | 9.1575 | Lys: 302 (17.179%) | Ser: 266 (15.131%) | Asp: 208 (11.832%) | Glu: 195 (11.092%) | Asn: 123 (6.997%) | SSSSSSSSSSSSRSSSSSTYTGSHDDSSEE |
| Sericin 3 | NP_001108116.1 | -1.4651 | 1271 | 123298.08 | 5.7076 | Ser: 554 (43.588%) | Gly: 152 (11.959%) | Gln: 94 (7.396%) | Asn: 85 (6.688%) | Lys: 79 (6.216%) | QSKASSSFSASSASESSSLSSDVNFEEKTD |
| Sericin 4 | BGIBMGA011896 | -0.9876 | 2270 | 251440.57 | 6.7156 | Lys: 223 (9.824%) | Thr: 218 (9.604%) | Ser: 215 (9.471%) | Glu: 205 (9.031%) | Gly: 169 (7.445%) | IHGGENTEAGGHSPGLVGGLFKYLLGGKTQ |
| Sericin 5 | KWMTBOMO06271 | -2.0155 | 1976 | 218868.3 | 9.6116 | Lys: 399 (20.192%) | Asp: 254 (12.854%) | Glu: 211 (10.678%) | Ser: 203 (10.273%) | Gly: 202 (10.223%) | SESASSSRSSYHSSSMQRNQKTSFDDDQNE |
| Sericin 6 / P150 | Sericin 6 / P150 | -0.6718 | 4552 | 467241.21 | 3.8151 | Thr: 1243 (27.307%) | Ser: 658 (14.455%) | Ala: 547 (12.017%) | Glu: 479 (10.523%) | Val: 297 (6.525%) | SKYGRCYCSCDLNSKPVFISLDGSALKPRR |
| Mucin-12 | XP_021205679.2 | -1.4977 | 3173 | 368381.36 | 4.137 | Thr: 583 (18.374%) | Glu: 522 (16.451%) | Gln: 478 (15.065%) | Pro: 218 (6.870%) | Leu: 202 (6.366%) | QGQNCFCTCDNTSAPKLIPLKDLPAGITGS |

**Supplementary Table S6.** Properties of P150 proteins in *B. mori* , *E. kuehniella* and *G. mellonella* . GRAVY: grand average of hydropathy; M. w.: molecular weight; pI: isoelectric point.

| Accession | Species | Family | Superfamily | GRAVY | Residue | M. w. (Da) | pI | Top1 AA | Top2 AA | Top3 AA | Top4 AA | Top5 AA |
|---|---|---|---|---|---|---|---|---|---|---|---|---|
| P150 | *Bombyx mori* | Bombycidae | Bombycoidea | -0.6718 | 4552 | 467241.21 | 3.8151 | Thr: 1243 (27.307%) | Ser: 658 (14.455%) | Ala: 547 (12.017%) | Glu: 479 (10.523%) | Val: 297 (6.525%) |
| WDD44665.1 | *Ephestia kuehniella* | Pyralidae | Pyraloidea | -0.8492 | 1741 | 173261.02 | 4.212 | Ser: 603 (34.635%) | Gln: 193 (11.086%) | Gly: 160 (9.190%) | Pro: 154 (8.845%) | Ile: 94 (5.399%) |
| XP_026763958.2 | *Galleria mellonella* | Pyralidae | Pyraloidea | -1.2812 | 1465 | 160622.85 | 4.7732 | Ser: 334 (22.799%) | Gln: 251 (17.133%) | Asn: 116 (7.918%) | Glu: 101 (6.894%) | Thr: 101 (6.894%) |
